# Honeybees vary communication and collective decision making across landscapes

**DOI:** 10.1101/2022.08.30.505816

**Authors:** Joseph Palmer, Ash E. Samuelson, Richard J. Gill, Ellouise Leadbeater, Vincent A.A. Jansen

## Abstract

Honeybee (*Apis mellifera*) colony foraging decisions arise from the waggle dances of individual foragers, processed and filtered through a series of feedback loops that produce emergent collective behaviour. This process is an example of animal communication at the height of eusociality, yet a growing body of evidence suggests that its value for colony foraging success is heavily dependent on local ecology. Although colonies are thought to vary their use of the waggle dance in response to local ecological conditions, this is yet to be empirically established. Here, we quantify waggle dance use based on colony level dance-decoding and show that the impact of dance use on collective foraging is clear in some colonies but nearly negligible in others. We outline how these estimates of dance use can be combined with land-use data to explore the landscape characteristics that drive collective foraging. Our methodology provides a means to quantify the real-world importance of a celebrated example of animal communication and opens the door to the exploration of the selection pressures that may have driven the evolution of this remarkable collective behaviour.

## Introduction

In group living animals, decisions can be taken collectively by integrating information from multiple individuals to produce a group behaviour that extends beyond that of the individual units (1). Such systems are self-organised, relying on the use of simple behavioural ‘rules’ that filter social information to produce an emergent collective outcome (2, 3). In honeybees, and other eusocial insects, the behavioural architectures that produce these emergent behaviours have become particularly complex. Honeybee colony foraging is coordinated via the waggle dances of individual foragers, which communicate food source locations to nestmates. A series of rules that determine when and how much bees dance mean that choices between feeding sites are made by the group (4–6). For example, because the number of dance circuits performed by a bee on returning from a food source reflects the net energetic benefits of the trip, more of the colony’s workforce will be recruited to the closer of two equally rich sources (7) or the richer of two equidistant sources (5). This filtered recruitment mechanism allows a colony to collectively allocate foraging effort adaptively without the need for any individual to compare resources.

Despite the complexity of this system, theory suggests that the value of dance recruitment for colony foraging success is not universal, but instead should vary significantly across ecological contexts (8–10). These predictions arise through the expectation that the likelihood of success of individual search varies according to the spatiotemporal distribution of food, such that waiting inside the hive to receive information is not always justified (9). Environments in which recruitment pays off are intriguing, because they are likely to have been important in driving the evolution of the waggle dance, yet empirical attempts to identify them have produced mixed results (11–18). Initial work, in which dances were disrupted and rendered meaningless by orientating the hive horizontally and thus removing the vertical axis that allows dancing bees to reference the sun’s position, tentatively linked the benefits of collective foraging to landscape heterogeneity (18). However, empirical attempts to systematically test this hypothesis have failed to provide support (11, 12), and dance disruption in challenging environments has sometimes even been associated with higher, rather than lower, foraging success (16). Consequently, no clear pattern has yet emerged with respect to the ecological conditions that shape the use of the dance communication system, (19), and which may have influenced its evolution.

Here, we set out an alternative approach to the question of when recruitment communication enhances foraging efficiency by identifying the ecological contexts in which honeybee colony foraging patterns are driven by recruitment. Bees do not always choose to dance on return from a foraging trip (20, 21) nor do they always seek out dances before leaving the hive (22). Even when they do follow dances, bees often ignore the spatial information in them when looking for forage sites (23). We hypothesise that this variation in individual use of the waggle dance will scale to the colony level, such that a colonies’ foraging behaviour will be described by differing extents of collective foraging through recruitment. To capture these dance use signatures, we develop a modelling approach to predict the distribution of foraging distances reported by individuals foraging entirely independently and compare it to the predictions of a model describing the distribution when a colony forages through a combination of independent search and waggle dance recruitment. Comparing these predictions to distances reported on the dancefloors of 20 real-world hives reveals that some colonies forage almost entirely collectively, whilst others are dominated by individual search. This provides a powerful inferential method to assess waggle dance use in naturally foraging colonies. Using the predictions of our model, we then explore how this variation in the impact of waggle dance communication could be evaluated from the perspective of landscape structure, demonstrating how our methodology could be used to assess the ecological circumstances under which collective foraging is important, and thus may have been significant in driving dance evolution. Put simply, by identifying those environments in which dances impact the shape of colony foraging patterns we can better understand why it evolved.

## Results

To establish how the distribution of foraging locations reported on the dancefloor might differ between colonies that rely on individual search and those that rely, to varying extents, on recruitment through dance communication, we simulated honeybees foraging in a landscape where resource patches were randomly placed in the environment. A forager could locate these under two different strategies: either acting as a scout and locating resources herself, or following a recruit strategy based on a random dance within the colony (6) (Fig. 1A, B, details in Materials and Methods). Note that “scouts” and “recruits” are not fixed behavioural categories, but rather, strategies adopted for a particular trip. As it is known in the simulation what proportion of individuals in the hive forage under what strategy, we can compare the distributions of foraging distances that should be visited by each type of forager.

**Figure 1.**
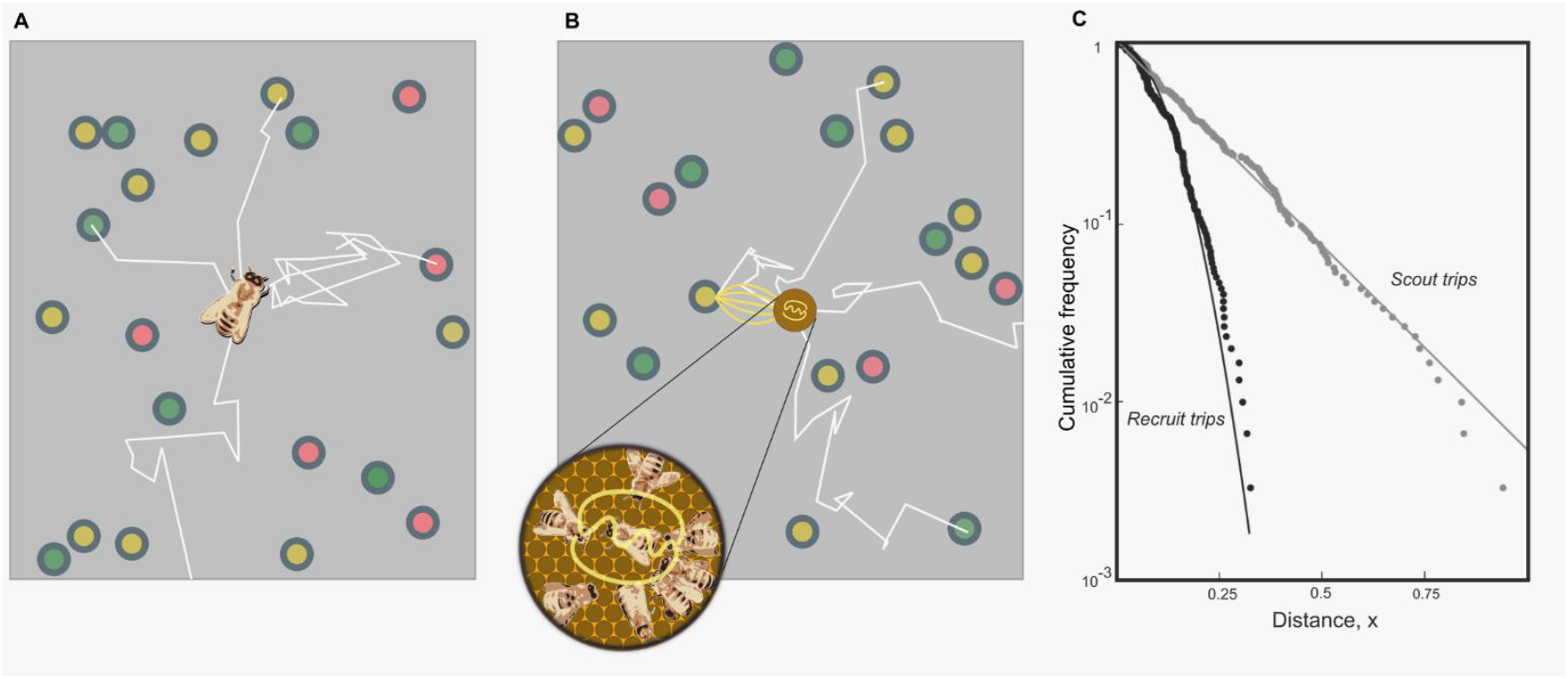
Simulating honeybee foraging. In our simulation model with scouting only (A), foragers leave the hive on a search path (white lines) and continue until they encounter a resource (circles, colours indicate different resource quality). When foraging with recruitment (B) foragers continue to identify resources in scouting trips (white lines) and convey this information on the dance floor (brown disc) where foragers can sample dances reporting on scouting and recruiting trips and follow these directions (yellow lines). (C) Complementary cumulative frequencies of foraging distances reported from scouting and recruit trips. Note the difference in the shape of the distributions. The scout distribution is best fit by an exponential (grey fit line), the recruit distribution by a Rayleigh distribution (black line).

The shapes of the resource distance distributions for bees engaging in the two types of foraging trips are different (Fig. 1C). The distance distribution for the scout trips is close to that of an exponential distribution (Fig. 1C), which is the nearest neighbour distance distribution for foragers operating in a one-dimensional environment (see Materials and Methods). Because scouts bias their dances so that sites offering higher net energetic gains are over-represented on the dancefloor (6), recruitment is more likely for closer sites (7, 20), assuming that food distribution is unbiased with respect to distance from the hive. The distribution of the distances reported for recruit trips thus differs from that of scouts and is captured by a Rayleigh distribution (Fig. 1C) which is the nearest-neighbour distribution in a two-dimensional environment (24) (see Materials and Methods). Combining these scout and recruit distributions in varying proportions in a mathematical model thus allows us to predict the distribution of waggle runs reported on the dancefloor of honeybee colonies under varying levels of recruitment (Fig 2, see Materials and Methods for details).

**Figure 2.**
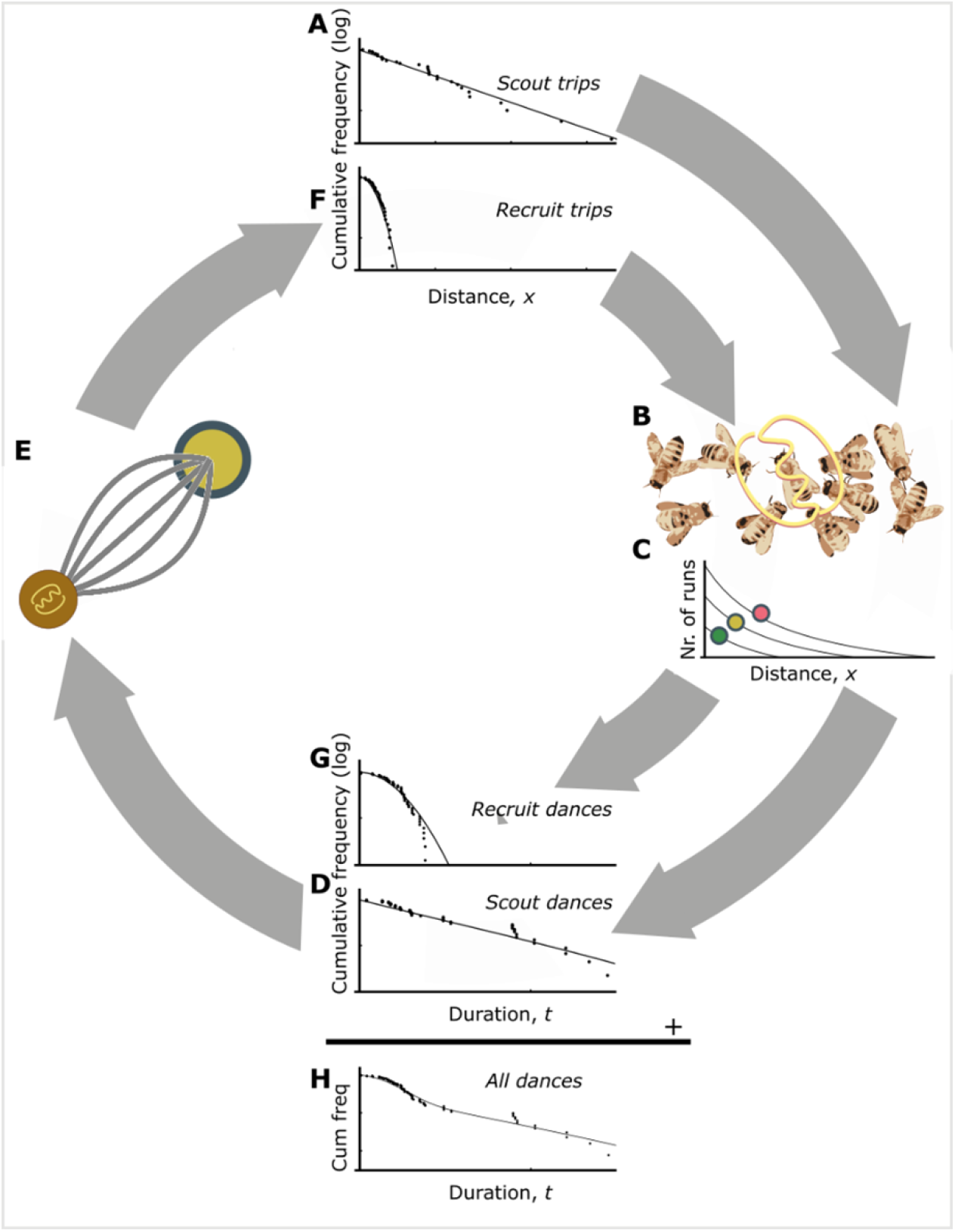
The use of communicating resource distances in honeybee foraging. The distances of resources encountered by scouts are distributed exponentially (A). These distances are communicated on the dance floor (B). Dances for resources that are closer or higher in quality are repeated more often (C). Thus, dances for more profitable resource are over-represented and sampling foragers are biased to the more profitable, closer resources (D). After successfully visiting advertised resources, recruits also dance for them leading to further amplification of this bias towards the most profitable resource in the vicinity of the hive (E). The distances of recruiting trips are than distributed through a Rayleigh distribution (F). Recruits also report the locations on the dance floor (B) and repeat their runs more often depending on the profitability of the location (C), leading to a distribution of durations of recruit dances (G). By taking together the dance distributions for scouts (D) and recruits (G) the distributions of all dances on the dancefloor can be found. The distances reported on the dance floor this are a mixture of the scout and recruiting trips and can be calculated from the distance distributions of the scouting and recruiting trips, taking the reporting bias into account (see Materials and Methods for detail).

To infer the use of communication and individual search in honeybee colonies foraging in ‘natural’ landscapes, we fitted our model to a pre-existing dataset (25) of 2827 decoded waggle dances from 20 observation hives at different locations (10 in central urban sites, 10 in agri-rural areas) in Southeast England, where dances had been recorded every fortnight from April to September 2017 (see Materials and Methods, Fig. 3A). For each site, we fitted the distances indicated in the waggle runs to two versions of our model: an “individual” model, where all forage sites are found through scouting, and a “collective” model, where the proportion of scout trips (*p*) can take on any value between 0 and 1. We used model selection (26) to determine which provided the better explanation of the data, and, if the collective model provided a better fit, to quantify the relative importance of communication through estimating the parameter *p*. In each case, we calculated the goodness-of-fit using a Kolmogorov-Smirnov (KS) test to ascertain if the model provided a plausible explanation of the data (27, 28).

**Figure 3.**
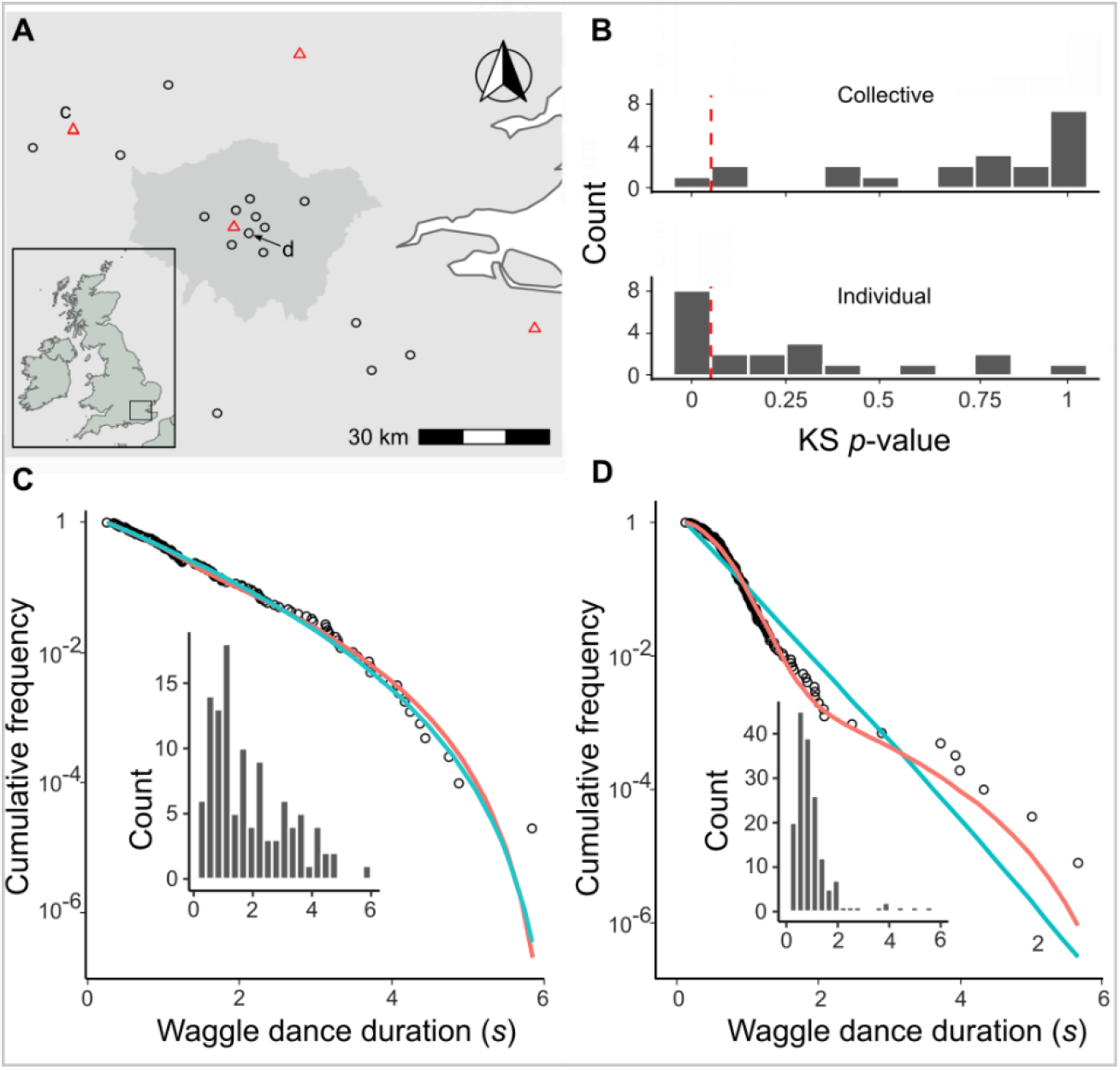
The honeybee foraging model fitted to data from 20 hives. (A) Location of study hives in Southern England, shaded area in the main plot indicates Greater London. For 16 hives for the collective foraging model provided best explanation (black circles) or for 4 hives the individual search model provided the best explanation (red triangles) as indicated by lowest AIC score. (B). Distribution of goodness of fit confidence values for each model fit to waggle run durations from each site. The p-value is derived from a bootstrapped two-sided KS test comparing the fitted model predictions to the empirical data, the red dashed line marks the significance threshold of 0.05. For values exceeding the threshold there is no statistically significant difference between the model and the data, indicating the model provides a good fit. For the hive in (C) the individual model (blue line) provided a better fit than the collective foraging model (red line). For the hive in (D) the collective foraging model (red line) provided a better fit than the individual model (blue line). The typical “hump” in the distribution in (D) which is indicative of contribution of recruitment dances (compare to Fig 3H). Panels show the compliment cumulative frequencies with binned frequency distributions as inset.

For 16 out of 20 study hives, the collective model provided a better description of the data than the individual model (Fig. 3A; Fig. S1). For the other 4 hives, despite the collective model having the higher maximum likelihood, the individual model had a lower AIC value and so is a more parsimonious description (Supplementary Table 1). In all but one hive, the collective model had a good fit (using a Kolmogorov-Smirnov statistic of *p* > 0.05, see Materials and Methods) to the empirical waggle run durations (Fig. 3B), whereas the individual model was significantly different to the observed data in 8 hives (Kolmogorov-Smirnov statistic *p* < 0.05, Fig. 3b). The hives shown in Figs 3C-D are representative examples showing the model fits where the individual (Fig. 3C) and collective (Fig. 3D) models fit best; for the full set see Fig S1. Note the closeness of the fit to the data, illustrating the overall quality of the model description.

Our model of individual foraging provides a more parsimonious description of foraging than a model of collective foraging in 4 hives. These results indicate that, whilst colony-level foraging is mostly comprised of a mixture of scout and recruit foraging trips, in some circumstances, it can be better described by individual foraging alone. Thus, in some environments, most foraging trips involve sites found through individual search rather than recruitment through dances. Note that this does not imply that these bees do not engage in dance following, because bees regularly follow dances but choose not to visit the advertised site (29), so it may be the case that dances are followed and ignored, not followed, or not performed. Nevertheless, although the mechanism by which flexibility is achieved at the individual level remains unclear, we have demonstrated that colonies themselves do not always forage collectively.

Our approach is not limited to a binary explanation of colony-level foraging as collective or independent. Quantifying the use of waggle-dance recruitment within colonies, as a proportion of all foraging trips, can be achieved by extracting the estimated proportion of recruit trips, 1 − *p*, henceforth termed “waggle dance use”. Variation in this metric can be used to identify contexts in which honeybees forage collectively, such as by examining its relationship with food distribution patterns (*e*.*g*. abundance/diversity/patchiness). Although our dataset was not specifically selected to represent variation in resource availability, because waggle dance use varied across our sites, our results nonetheless illustrate conclusively that waggle dance use varies with landscape structure.

Our hives were located at either urban sites, where forage for social bees is typically relatively abundant, diverse and consistent across the season, or agricultural-rural (agri-rural) sites, which are thought to be characterized by long periods of relatively sparse forage interspersed with temporary high abundance through mass-flowering crops (30). Accordingly, while waggle dance use was typically high for both land-use types, agricultural sites showed considerable variation (Fig. 4A), which was less apparent within the urban sites. Consequently, this variation provides an opportunity to demonstrate how our models’ predictions might correlate with land-use. To assess this, we performed a GIS-based land-use classification to obtain a quantitative land-use profile for each of the agri-rural sites (31), and used a Partial Least Squares analysis (32) to identify the combinations of land-use types which most explained the variation in our estimates of dance use. A single principal component explaining ~73% of the variation in waggle dance use correlated positively with arable land cover (29% of total land area; Supplementary Table S2) and negatively with built-up area (17% of land area; Fig. 4C; beta regression: *R*^2^ = 0.73, *ϕ* = 4.9, P < 0.05, Fig. 4B). This suggests that forage was more likely to be found through recruitment in relatively built-up areas within the agri-rural landscape.

**Figure 4.**
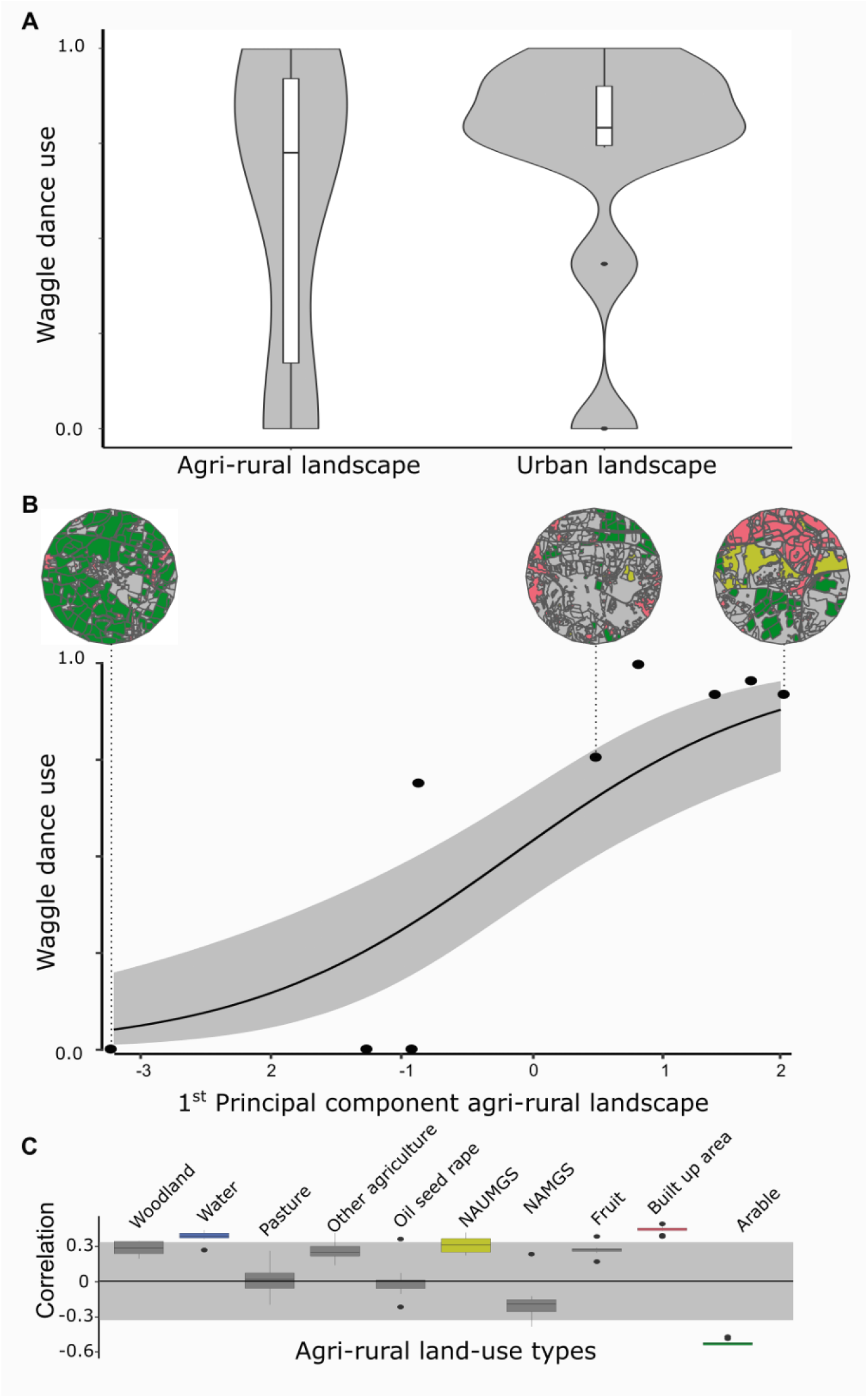
Collective foraging correlates with land-use. Within the agricultural land-use category, we found considerably variation in waggle dance use across sites (A) and we thus sought to explore whether this could be explained by differences in land-use across sites using PLS analysis (B) Beta regression shows the relationship (black line) between first principal component and waggle dance use, with 95% CI shown by the grey shaded area. Boxplots in panel (C) illustrate the range of correlations between the first principal component and each land-use type (colours correspond to the land use as shown in circular inset maps for selected sites), following jacknife resampling. Correlations outside the shaded area significantly contribute to the first principal component. NAUMGS (resp. NAMGS) stands for non-agricultural unmanaged (resp. managed) green space.

## Discussion

Our results provide empirical evidence that there is considerable variation in the impact of waggle dance recruitment on colony behaviour in naturally foraging colonies, and therefore, in the extent to which communication drives collective foraging by honeybee colonies. The foraging patterns of some colonies bear a clear hallmark of recruitment, while others do not. Previous work has sought to illustrate the impact of dance communication for colony foraging by preventing communication and evaluating the consequences for food intake, relative to control colonies (11– 18). Our approach is distinct and found dancing was unrestricted in all colonies, but under some circumstances had no detectable impact on the spatial allocation of colony foraging effort between resources, producing foraging patterns that were indistinguishable from those that would have occurred through independent search alone.

We established this through a novel statistical inferential methodology that enabled us to quantify the proportion of a colony’s foraging that is driven by collective or individual search. Our model distinguishes between sites found through individual search and through recruitment by assuming the existence of a “filter” that leads to sites that are closer to the hive being over-represented on the dancefloor. This filter arises through two potential mechanisms that are inherent properties of the dance communication system. Firstly, more distant forage sites take longer to reach, and so the number of hive visits per unit time reduces accordingly, such that the opportunities to dance are more limited (7). Secondly, it is well established that the number of dance circuits performed on a forager’s return to the hive is determined by the energetic efficiency of the trip (4). Assuming unbiased distribution of nectar concentration with respect to the hive’s location, closer sites will therefore elicit both more dances, and more dance circuits per dance (sometimes interpreted as “livelier dances”; but see (6)), and each single dance circuit increases the likelihood that a follower will be recruited to a resource (33). These filtering properties produce the key differences in the distribution of distances reported by scouts and recruits that are identified by our approach. When these different distributions were combined, they produced a composite distribution with a characteristic long tail of distant foraging sites visited almost entirely by scouts. Our predictions are based on a simplified description of the recruitment process, without attempting to capture its full complexity, but nonetheless achieve a striking fit to data from real-world hives.

What mechanism underlies this variation in the impact of waggle dance communication on colony forage site choice? It seems unlikely that dance-followers in different colonies followed different rules about how to respond to dances (but see (16)). Rather, the same rules may have produced different outcomes at the colony level depending on resource availability patterns. Despite its well-known role in recruiting bees to newly discovered food sources (20, 29, 33), the majority of dances within hives indicate forage sites that are already known to the dance-followers (22), serving to motivate continued foraging (33). In colonies where individual search dominated forage patterns, we expect that bees still engaged in dance-following behaviour but were rarely recruited to new sites through it. Potentially, when individuals’ known patches became depleted, fewer dances indicating novel alternatives were available because such sites did not exist, or because they were not of high enough quality to elicit many dance circuits. Rather than wait for information, such foragers might then seek out sites through individual search (9, 16). This possibility invites exploration of the impact of forage availability and heterogeneity on resource representation on the dancefloor.

We found that waggle dance use varied with landscape structure. Our hive locations derived from a separate study (25) and were not specifically chosen to capture variation in food distribution, but they spanned land-use types that follow established general patterns in abundance and diversity. Agricultural land in the UK is typically considered nutritionally poor for bees, with large areas of limited food availability punctuated by brief availability of rich mass-flowering crops in some areas, while more urbanized residential areas that contain gardens are relatively forage rich and less variable across the season (30, 34, 35). Accordingly, we found high variation in waggle dance use in the agri-rural category, and that variation could be mapped to land-use. Specifically, waggle dance use decreased as built-up areas gave way to arable land. Our dataset comprised recordings across an entire foraging season and so lacks the temporal resolution to directly link this result to food availability, which would have varied across the season, but our findings suggest that the impact of waggle dance use on colony foraging may be more pronounced when forage is rich and abundant. On this basis, further work could focus on rural locations systematically chosen to represent a range of floral diversity and abundance (e.g. (36)), with the temporal resolution to focus on specific periods of the year, to identify those ecological contexts in which dance communication has a detectable impact on colony foraging.

This is important because although it is widely assumed that some aspect of the tropical forests in which *Apis* evolved favoured the evolution of the dance (19). Current attempts to identify those critical features rely upon labour-intensive dance-disorientation protocols, whereby dance communication is actively prevented and the consequences for foraging efficiency monitored (13, 14, 18, 36, 37). Such studies are elegant in design but logistically challenging to perform at a scale that allows inter-colony variation in foraging environments at sufficient replication (but see ((36)), particularly given that multiple landscape variables may interact to determine the utility of dance communication. Given the recent development of automated dance-decoding protocols (38, 39), now also validated for field-based videos (40), our methods provide a time and labour efficient means to identify the ecological and evolutionary environments in which this iconic example of animal communication shapes collective behaviour. We have developed a means to establish when dancing leaves a meaningful imprint on colony foraging patterns, narrowing down the search for ecological contexts in which it significantly improves colony fitness.

## Materials and Methods

### Overview

To predict the distribution of forage site distances reported by bees employing individual search (scouts) or following dances (recruits), we first simulated the foraging behaviour expected under each strategy. We then used the resulting distributions to inform a collective model of colony behaviour in which the proportion of bees performing scout trips (*p*) can take any value between 0 and 1. Using a pre-existing dataset of foraging distributions extracted from videos of waggle dances in 20 hives, for each hive we compared the fit of this collective model with a model in which foraging is entirely independent. Finally, for each hive within the agri-rural category (n=10 hives), we calculated a multivariate land-use profile to describe the surrounding landscape and performed Partial Least Squares analysis to establish how the proportion of bees acting as recruits (“waggle dance use”, estimated from the collective model) varied with land use.

### Simulation

A circular foraging environment was created with radius *r* = 2.5. The number of resources in the environment was generated as a random Poisson variable with a mean of 1/5000 multiplied by the area of the environment. Resources are located at polar coordinates with a uniformly selected angle, *θ*, between 0 and 2*π* and a radial value, *ρ*, between 0 and *r*, determined by the square root of the uniform position values multiplied by *r*. Each location was assigned to an instance of a resource object, where resource quality was randomly allocated between 0 and 10. Resource profitability is a product of this quality and the distance of the resource to the centrally located hive.

One hundred honeybee objects forage in this environment. Scouts follow a random path through the environment generated by sampling a uniform random step length and angle. The number of paths the scout draws when searching is also determined as a uniform random number. Each straight line in the random path is converted to a rectangle with length equal to the path section length and a constant width of ~0.01 to represent an area the scout searches along that path. Of all the resources contained in the boxes drawn from the scout’s path, the one closest to the colony is selected as the resource patch that the scout will report if the quality of the resource exceeds a minimum threshold. Communication is simulated by pooling all the resource patches found. If no resources are contained in the scout’s path, no resources will be added to the scout pool.

Recruits are honeybee objects that do not perform individual searches for forage, but instead sample from the pool of resource objects reported by scouts. The probability of sampling a resource is skewed by its profitability, mimicking the profitability bias known to occur in the dances of real-world scouts (6). Recruits then visit these resources, and in the next iteration will add their resource to the pool of scout dances. Consequently, the pool of dances represents resources discovered by scouts and resources exploited by recruits. When a resource is depleted, it is removed from the environment and so any foragers that had been visiting it must select a different resource from the dance floor.

The simulation was run for 100 iterations, in which all distances reported by scouts and recruits were recorded and combined every 5 time steps. We fit an exponential and minimum of an exponential distribution to the distribution of foraging distances reported by the (Fig 1. A) scout and (Fig 1. B) recruit objects, by deriving the maximum likelihood estimate for each model fit on each data source through their analytical solutions: 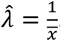, minimum of the exponential with a minimum foraging distance: 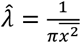. As the exponential assumes that distributions start from 0, the data were transformed by subtracting the minimum foraging distance from all foraging distances (*x* = *x* − *min*(*x*)) before fitting.

All simulation code was written in Python version 3.9 and uses the Pandas (41) and Scipy (42) packages.

### Mathematical Model

The duration of the waggle run of a dance circuit represents the distance flown by the bee, and the two are linearly related (43, 44). To describe the distribution of waggle run durations on the dance floor we formulated a generic model for the duration of waggle dances, in which resource patches are randomly placed in the environment. Foragers scout for these patches. Upon visiting a resource patch, foragers translate the profitability of a resource into the number of repeats of the dance, also called “dance circuits” (Fig 2). The number of dance circuits is a function of quality and distance. Recruits sample random dances and report the location of successful visits to resource patches on the dance floor. Through the feedback and over-representation of profitable resources on the dance floor, recruits will converge to visiting the most profitable resource in vicinity of the hive. The distribution of waggle run durations is the superposition of scouting and recruiting trips.

The distance after which the first resource is discovered approximately follows an exponential distribution (given by λ*e*^−λ*x*^). Through the feedback mechanism that the dance floor provides, the colony can, collectively, locate the most profitable resource in its environment. For randomly placed resources in a two dimensional environment the distance to the nearest point is distributed according to a Rayleigh distribution (given by 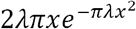) (24). Our simulation model shows that this describes the distances at which recruits visit resources well. Knowing the distance distributions of scout and recruit trips we then assume that the proportion *p* of all trips are scout trips. With this information we can specify the distributions of distances on the dance floor (see Supplementary Material for details).

We implemented this in a full model that describes the distance distribution of an environment that has *n* different resource types (See Supplementary Material). In the full model the number of parameters increases with 2*n* (each resource needs a parameter for the scout and recruit distribution). Even if the number of resources is low, the model tends to overfit. To facilitate estimation of the parameter *p* we therefore used a simplified model to estimate the fraction of scout trips, where *m* represents the lowest duration considered, and here a minimum waggle run duration in the data set.

In the simplified the model we assumed that the number of dances depends weakly on distance and there is a sizable quality differences between resources of a non-negligible size and that there is a sizable intensity of the high-quality resource (See Supplementary Material for detailed derivation). Foragers on scouting trips are more likely to report larger distances than foragers on recruiting trips. By linearising the function that translates the profitability into the number of waggle dance run for the largest waggle run duration and normalising, we arrive at simplified distribution for waggle run durations for scouting trips:

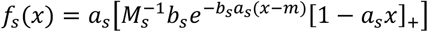

where we used the shorthand *x*_+_ = max(*x*, 0) The parameter 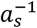 is the maximum waggle run duration by scouts, *a*_*s*_*b*_*s*_ is the intensity of resources found by scouts and 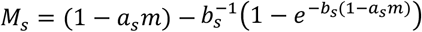 is factor that normalises the distribution.

Recruit trips will be predominantly to high-quality resources. Only if the nearest high-quality patch is very far away will there be a more profitable patch of lesser quality available, and this happens only rarely if the intensity of the best quality resource is sizable. After linearising the function that translates the profitability into the number of waggle dance runs for short waggle run durations and normalising the distribution of waggle run durations reported from recruit trips in the simplified model is:

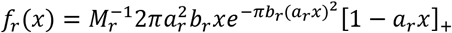

where

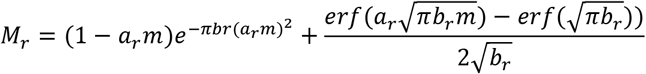

is the normalisation factor, the parameter *a*_*r*_ is the rate with which dances repeats depends on distance for recruit trips and 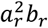 is the intensity of high-quality resources reported by recruited foragers.

The simplified distribution function is

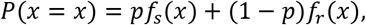

which we used for parameter estimation and model fittings.

### Data collection

Details of data collection, waggle dance decoding and classification of land-use types can be found in full in the Materials and Methods section of (25). Briefly, observation hives were located at apiaries in either central London (UK) or the surrounding agricultural land and were each located at least 2km apart. Visits took place every fortnight between April and September 2017. On each visit, two hours of continuous waggle dance data was recorded by training a camcorder onto the dance floor. The footage of the dances was decoded manually following methods in (45).

### Model fitting

For each real-world hive from the observed dataset, we fitted the distances indicated in the waggle runs to two versions of our model: an “individual” model, where all forage sites are found through scouting, and a “collective” model, where the proportion of scout trips (*p*) can take on any value between 0 and 1.

All models were fit using Maximum likelihood estimation (26) by summation of the log of the simplified distribution function outlined in the methods section: model. The numerical optimisation routine is written in c++ and uses the Nelder-Mead simplex algorithm (46) implemented in the ‘NLopt’ library (47) and interfaced to R using ‘Rcpp’ (48).

The most parsimonious model was identified using Akaike information criterion (AIC) and Akaike weights (26, 49). Goodness of fit was assessed using the two-sample Kolmogorov-Smirnov (KS) test (28) and implemented in R using the ks.boot function of the package ‘Matching’ in R (50).

All analysis code was written in R (51).

### Partial Least Squares analysis

For each hive, our modelling process resulted in an estimated proportion of recruit trips, 1 − *p*, henceforth termed “waggle dance use”. This proportion was highly variable within the agri-rural category, and so within this group, we sought to identify land-use variables that covary with waggle dance use. For each site, we performed Partial Least Squares analysis based on proportional coverage within 10 land-use categories within a 2.5km radius around each hive (Table 1, see (25) for full classification methods). Prior to conducing the PLS we removed any sites in for which both individual and collective model fit was poor (n=1 site).

As our estimates of waggle dance use are continuous on the interval [0,1] we used the R package plsRbeta (52) to conduct the PLS and performed a beta regression on the results using the R package betareg (53). As the betareg package only works on the open interval (0,1) the data, *x*, was transformed using the following equation: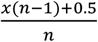 as outlined in the betareg package documentation. After analysis the data was back-transformed to the original values for the plots in Fig 4.

To test robustness, we performed jackknifed resampling by removing each site in turn before re-rerunning the PLS analysis, recoding the loadings for each iteration (see Supplementary Material for loadings with each site removed). The PLS loadings for each land-use type are plotted as a box plot in Fig 4. to show the spread of these variable types. A loading was determined to be significantly correlating with the first principal component if contributed more than its expected variance.

Note that significant correlations with land-use types that represent very small proportions of total land cover were not interpreted further (unmanaged green space; Supplementary Table S2; Fig. 4C. Unmanaged green space was also not supported by jackknife analysis).

## Acknowledgments

We are grateful to the beekeepers who allowed installation of observation hives at their apiaries for collection of the original dataset.

## Data and materials availability

All the data and code required to reproduce this article can be found at: https://doi.org/10.5281/zenodo.7025591

## Author Contributions

Designed study: JP, RG, EL, VJ. Performed research: JP, AS, EL, contributed new reagents or analytic tools: JP, VJ, EL. Analysed data: JP, AS. Wrote the paper: Original Draft: JP, VJ, EL Writing - review & editing: JP, RG, EL, VJ. Supervision: RG, EL, VJ.

