## Supplementary material for "Honeybees vary communication and collective decision making across landscapes"

### The derivation of the full model for the distribution of distances on the dance floor.

We will build a simple model to describe the distribution of distances reported on the dance floor. We assume that there are $n$ different resources and they are distributed according to a spatial Poisson point process. The Poisson point process places food sources randomly in the environment of the hive with intensity $\lambda_i$ for resource $i$. The quality of that resource we will denote $q_i$. For convenience, we will order the resources $q_i$ from small to large such that $q_j \geq q_i$ if $j > i$, and $q_n$ is the resource with the highest quality, which we assume is always present in the environment, hence $\lambda_n > 0$, and has a resource quality exceeding unity, hence $q_n > 1$. Under this ordering, we define $m$, for which $1 \leq m \leq n$, as the lowest index for which $q_m > 1$ hence, $q_i < 1$ for all $i = 1, \ldots, m-1$

**The distribution of scout dances.** As a scout travels a small distance $\mathrm{d}x$ its search will cover an area $c_s\,\mathrm{d}x$. The constant $c_s$ depends on width of the area scanned, the detection rate and the degree to which a the forager's path covers new ground. If we incorporate these factors into the constant $c_s$ the intensity of discovering a resource over a path of distance $\mathrm{d}x$ is $c_s\,\mathrm{d}x$. The probability density of $x$, the distance travelled after which the first source is discovered follows a distribution

$$\lambda_s e^{-\lambda_s x}$$

where $\lambda_s = c_s \sum_{i=1}^{n} \lambda_i$.

The profitability, which depends on quality and distance, of the source dictates the number of repeats of the dance. What is the distribution of dances on the dance floor? Let's assume that there is a function that translates the profitability into the expected number of runs of the waggle dance, $\phi(q_i, x)$, which is a function of quality an distance. We assume that if this function has a non-zero value it decreases with distance, and increases with quality, hence $\frac{\partial \phi(q,x)}{\partial q} > 0$ and $\frac{\partial \phi(q,x)}{\partial x} < 0$ if $\phi(q,x) > 0$. There is a minimum quality level of the resource that elicits dancing, and we have set this level to unity, such that if $q \leq 1$ then $\phi(q,x) = 0$ for $x \geq 0$. It follows that for each $q_i > 1$ there is a maximum distance $x_{i,\max} > 0$ for which bees do not dance, that is $\phi(q_i, x) = 0$ for $x \geq x_{i,\max}$.

The expected number of dances for a resource of type $i$ located at distance $x$ is $c_s \lambda_i \phi(q_i, x) e^{-\lambda_s x}$. The expected total number of scout waggle dances on the dance floor is given by

$$M_s = \sum_{i=m}^{n} \int_0^{\infty} \frac{c_s \lambda_i}{\lambda_s} \phi(q_i, x) \lambda_s e^{-\lambda_s x}\,\mathrm{d}x$$

and the distribution of scout dances is

$$\frac{\lambda_s e^{-\lambda_s x} \sum_{i=m}^{n} \frac{c_s \lambda_i}{\lambda_s} \phi(q_i, x)}{M_s}$$

If the bees go out, scout for resources and dance for them this, and there is no further processing of information, this would be the distribution one would expect.

**The distribution of recruit dances.** First, we consider a simple scenario and we assume that all resources are of equal quality $q$ with $q > 1$. The scouts that report back on the dance floor cover the surroundings of the hive in their forays. How well they cover, depends on the number of scouts, how well their collective paths cover the area and chance of discovery of an item. Let the fraction of sources that can be discovered by the collection of scouts be $c_r$. We assume $0 \leq c_r \leq 1$.

The chance of discovering food source within a radius of $x$ is $e^{-c_r \lambda_r \pi x^2}$, where $\lambda_r = \sum_{i=1}^{n} \lambda_i$ and therefore the chance of having the nearest food source that the recruits report is least a distance $x$ away is $P(\underline{x} \geq x) = e^{-c_r \lambda_r \pi x^2}$. The distribution of the nearest food source that recruits report is

$$2 c_r \lambda_r \pi x e^{-c_r \lambda_r \pi x^2}.$$

The distribution of recruit dances on the dance floor is:

$$\frac{\phi(q, x) 2 c_r \lambda_r \pi x e^{-\lambda_r \pi x^2}}{M_r}$$

where

$$M_r = \int_0^\infty \phi(q, x) 2 c_r \lambda_r \pi x e^{-c_r \lambda_r \pi x^2} \, \mathrm{d}x.$$

If the resources differ in quality, the same rationale can be applied. Consider two locations that differ in resource quality and distance, the hive can prefer the further location if it is sufficiently better. Likewise, it can prefer a lesser quality location if it is sufficiently nearer. Given that the hive has found a resource at distance $x$ of quality $q_i$ we define a threshold distance $\xi_{ij}(x)$ at which a resource of type $q_j$ has the same profitability. If $q_j \geq q_i$, and $0 \leq x \leq x_{i,\max}$ then $\xi_{ij}(x)$ satisfies

$$\begin{cases} \phi(q_i, x) = \phi(q_j, \xi_{ij}(x)) & \text{if } 0 \leq x < x_{i,\max} \\ \xi_{ij}(x_{i,\max}) = x_{j,\max} & \text{if } x = x_{i,\max} \end{cases}$$

If $q_j < q_i$, and $x < x_{i,\max}$ then $\xi_{ij}$ satisfies

$$\begin{cases} \xi_{ij}(x) = 0 & \text{if } 0 \leq x \leq \xi_{ji}(0) \\ \phi(q_i, x) = \phi(q_j, \xi_{ij}(x))) & \text{if } \xi_{ji}(0) < x_{i,\max} \\ \xi_{ij}(x_{i,\max}) = x_{j,\max} & \text{if } x = x_{i,\max} \end{cases}$$

This relation defines the threshold distance $\xi_{ij}(x)$ as a function of $x$ for $0 \leq x \leq x_{i,\max}$. It follows that $\xi_{ii}(x) = x$. Because $\phi(q_i, x)$ increases with $q$ if $\phi(q_i, x) > 0$ it follows that $\phi(q_i, x) > \phi(q_j, x)$ if $q_i > q_j$. As a consequence $\xi_{ij} < x$ if $i > j$ and $\xi_{ij} > x$ and if $i < j$.

The probability to find the nearest resource $i$ at distance $x$ is given by the Rayleigh distribution $2\pi c_r \lambda_i x e^{-\pi c_r \lambda_i x^2}$. For $x \leq x_{i,\max}$ the chance of there not being a source if type $j$ within a radius of $\xi_{ij}(x)$, which is $e^{-\pi c_r \lambda_j \xi_{ij}(x)^2}$. The probability that resource $i$ at location $x$, $0 \leq x \leq x_{i,\max}$ is the most profitable is therefore

$$P(\text{most profitable resource } i \text{ is at distance } x) = 2\pi c_r \lambda_i x e^{-\pi \sum_{j=1}^{n} \lambda_j c_r \xi_{ij}(x)^2}$$

and 0 if $x > x_{i,\max}$ and the probability of finding the most profitable location at distance $x$ is

$$P(\text{most profitable resource is at distance } x) = \sum_{\forall i : x_{i,\max} > x} 2\pi c_r \lambda_i x e^{-\pi \sum_{j=1}^{n} c_r \lambda_j \xi_{ij}(x)^2}.$$

For $x > x_{n,\max}$ the most profitable resource cannot be determined, as all have zero profitability.

The probability of encountering a recruit reporting a distance $x$ on the dance floor is

$$M_r^{-1} \sum_{i=m}^{n} \phi(q_i, x) \lambda_i 2\pi c_r x e^{-\pi \sum_{j=1}^{n} c_r \lambda_j \xi_{ij}(x)^2}$$

where $M_r$ is the normalisation factor

$$M_r = \int_0^\infty \sum_{i=m}^{n} \phi(q_i, x) \lambda_i 2\pi c_r x e^{-\pi \sum_{j=1}^{n} c_r \lambda_j \xi_{ij}(x)^2} \, \mathrm{d}x.$$

**The full model.** The combined distribution of distances of scout and recruit dances on the dance floor is given by

$$P(\underline{x} = x) = p \frac{\lambda_s e^{-\lambda_s x} \sum_{i=1}^{n} \frac{c_s \lambda_i}{\lambda_s} \phi(q_i, x)}{M_s} + (1-p) \frac{\sum_{i=1}^{n} \phi(q_i, x) 2\pi c_r \lambda_i x e^{-\pi \sum_{j=1}^{n} c_r \lambda_j \xi_{ij}(x)^2}}{M_r}.$$

This is a generic model for the distribution of distances reported on the dance floor.

### The simplified model

The full model can be used to calculate the likelihood for a given data set, which can be used to estimate parameters. The full model has $2n + 3 + k$ parameters: $\lambda_1 \ldots \lambda_n$, $q_1 \ldots q_n$, $c_s, c_r$ and $p$ plus $k$ parameters needed to specify the function $\phi$. Even though it is possible to reduce the number of parameters through a scaling, e.g. by defining $\lambda_i^r = c_r \lambda_i$ and $c = \frac{c_s}{c_r}$, but even if the number of resources is low, it turned out to be difficult to estimate the parameters even if with a fair sized data sets. To assist in the estimation of the key parameters we therefore formulated a simplified model.

We can simplify the model by assuming that the dependence of the number of dances depends less on the distance and more on quality. We assume that that dependence of the profitability assessment only weakly depends on distance and that there is quality differences between resources of a decent size, so that there are a large threshold distances at which resources have the same profitability. If that is the case then, if a resource is discovered at a small distance from the hive, resources if less quality than this one, are not worth dancing for, whereas there is a large distance over which superior resources can be detected that will have higher profitability. In mathematical terms, under this assumption, for small $x$, $\xi_{ij}(x) = 0$ if $j < i$, $\xi_{jj}(x) = x$ and $\xi_{ij}(x)$ is large if $j > i$. We thus have $\sum_{j=1}^{n} \lambda_j \xi_{ij}(x) = \sum_{j=i}^{n} \lambda_j \xi_{ij}(x)$ for $i < n$ and $\sum_{j=1}^{n} \lambda_j \xi_{nj}(x) = \lambda_n x$. Moreover, as the $\xi_{ij}s$ will be large, $\sum_{j=i}^{n} \lambda_j \xi_{ij}(x) \gg \lambda_n x$. We can approximate

$$\sum_{i=1}^{n} \phi(q_i, x) 2\pi c_r \lambda_i x e^{-\pi \sum_{j=1}^{n} c_r \lambda_j \xi_{ij}(x)^2} \approx \phi(q_n, x) 2\pi c_r \lambda_n x e^{-\pi c_r \lambda_n x^2}$$

The recruits on the dance floor are most likely to report short distances. By expanding the function $\phi(q_n, x)$ around $x = 0$ we get $\phi(q_n, x) = \left[ \phi(q_n, 0) + x \left. \frac{\partial \phi(q_n, x)}{\partial x} \right|_{x=0} + O(x^2) \right]_+$ we can approximate the recruit distribution as:

$$M_r^{-1} \sum_{i=m}^{n} \phi(q_i, x) \lambda_i 2\pi c_r x e^{-\pi \sum_{j=1}^{n} c_r \lambda_j \xi_{ij}(x)^2} \approx \frac{[1 - a_r x]_+ \, 2\pi a_r^2 b_r x e^{-\pi b_r (a_r x)^2}}{1 - \frac{\mathrm{erf}\left(\sqrt{\pi b_r}\right)}{2\sqrt{b_r}}}$$

with $a_r = -\phi(q_n,0)^{-1} \left.\frac{\partial\phi(q_n,x)}{\partial x}\right|_{x=0}$ and $b_r = c_r\gamma_n a_r^{-2}$ and where we used

$$\phi(q_n,0)\int_0^{a_r^{-1}} (1-a_r x)2\pi a_r^2 b_r x e^{-\pi b_r (a_r x)^2}\,\mathrm{d}x = \phi(q_n,0)\left(1-\frac{\operatorname{erf}\left(\sqrt{\pi b_r}\right)}{2\sqrt{b_r}}\right)$$

and cancelled the term $\phi(q_n,0)$.

The scouts on the dance floor are more likely to report large distances. For distances around the largest distance for which scouts dance, $x_{n,\max}$, the $\phi(q_i,x) = 0$ for all $i < n$. We therefore expand $\phi(q_n,x)$ around $x_{n,\max}$ to get $\phi(q_n,x) = \left[(x - x_{n,\max}) \left.\frac{\phi(q_n,x)}{\partial x}\right|_{x=x_{n,\max}} + O(x^2)\right]_+$. With this, we can approximate the scout distribution as:

$$\frac{\lambda_s e^{-\lambda_s x}\sum_{i=m}^{n}\frac{c_s\lambda_i}{\lambda_s}\phi(q_i,x)}{M_s} \approx \frac{a_s b_s e^{-b_s a_s x}\left[1-a_s x\right]_+}{1-b_s^{-1}\left(1-e^{-b_s}\right)}$$

with $a_s = x_{n,\max}^{-1}$ and $b_s = \gamma_s a_s^{-1}$ and where we used

$$\frac{c_s\lambda_i}{\lambda_s}\left.\frac{\phi(q_n,x)}{\partial x}\right|_{x=x_{n,\max}}\int_0^{a_s^{-1}} a_s b_s e^{-b_s a_s x}(1-a_s x)\,\mathrm{d}x = \frac{c_s\lambda_i}{\lambda_s}\left.\frac{\phi(q_n,x)}{\partial x}\right|_{x=x_{n,\max}}\left(1-b_s^{-1}\left(1-e^{-b_s}\right)\right)$$

and cancelled the term $\frac{c_s\lambda_i}{\lambda_s}\left.\frac{\phi(q_n,x)}{\partial x}\right|_{x=x_{n,\max}}$.

The distribution then simplifies to

$$P(\underline{x}=x) = p\frac{\left[1-a_s x\right]_+ a_s b_s e^{-b_s a_s x}}{1-b_s^{-1}\left(1-e^{-b_s}\right)} + (1-p)\frac{\left[1-a_r x\right]_+ 2\pi a_r^2 b_r x e^{-\pi b_r (a_r x)^2}}{1-\frac{\operatorname{erf}\left(\sqrt{\pi b_r}\right)}{2\sqrt{b_r}}}.$$

The simplified model has 5 parameters, and is therefore simpler than the full model for $n \geq 2$, which has $2+2n+k$ parameters.

### The simplified distribution with a minimum value

Let $m$ be the lowest distance that is recorded, for instance because there is a minimum duration of the waggle dance. We assume that $m < \xi_{ji}(0)$ for all $j < i$. For the simplified recruit distribution we now expanding the function $\phi(q_n,x)$ around $x = m$ to get $\phi(q_n,x) = \left[\phi(q_n,m) + (x-m)\left.\frac{\partial\phi(q_n,x)}{\partial x}\right|_{x=m} + O(x^2)\right]_+$. The simplified recruit distribution becomes:

$$\frac{\left[1-a_r x\right]_+ 2\pi a_r^2 b_r x e^{-\pi b_r (a_r x)^2}}{\left(1-a_r m\right)e^{-\pi b_r (a_r m)^2} + \frac{\operatorname{erf}\left(a_r\sqrt{\pi b_r}m\right)-\operatorname{erf}\left(\sqrt{\pi b_r}\right)}{2\sqrt{b_r}}}$$

with $a_r = m - \phi(q_n,m)^{-1}\left.\frac{\partial\phi(q_n,x)}{\partial x}\right|_{x=m}$ and $b_r = c_r\gamma_n a_r^{-2}$.

The scout distribution is only affected through the normalisation factor and becomes:

$$a_s\frac{b_s e^{-b_s a_s (x-m)}\left[1-a_s x\right]_+}{\left(1-a_s m\right) - b_s^{-1}\left(1-e^{-b_s(1-a_s m)}\right)}$$

The distribution, for $x \geq m$ then becomes

$$\begin{aligned} P(\underline{x} = x) &= p a_s \frac{b_s e^{-b_s a_s (x-m)} \left[1 - a_s x\right]_+}{(1 - a_s m) - b_s^{-1}\left(1 - e^{-b_s(1-a_s m)}\right)} \\ &\quad + (1-p) \frac{\left[1 - a_r x\right]_+ 2\pi a_r^2 b_r x e^{-\pi b_r (a_r x)^2}}{(1 - a_r m)\, e^{-\pi b_r (a_r m)^2} + \frac{\mathrm{erf}\left(a_r \sqrt{\pi b_r} m\right) - \mathrm{erf}\left(\sqrt{\pi b_r}\right)}{2\sqrt{b_r}}}. \end{aligned}$$

The same function describe the distribution of the dance durations, where dances have a minimum duration. This assumes a linear relation between duration and distance.

### Model fits to all sites

Table S1. Fit results for the fitting of the waggle dance model to each site.

| | site | model | loglikelihood | AIC | delta_AIC | rAIC | wAIC | p | bs | br | as | ar | k | ks_statistic | ks_pvalue |
|---|---|---|---|---|---|---|---|---|---|---|---|---|---|---|---|
| 1 | BEL | collective | -181.41 | 372.83 | 1.83 | 0.40 | 0.29 | 0.65 | 7.04 | 0.81 | 0.05 | 0.17 | 5 | 0.10 | 0.71 |
| 2 | BEL | individual | -183.50 | 371.00 | 0.00 | 1.00 | 0.71 | 1.00 | 4.50 | | 0.07 | | 2 | 0.09 | 0.79 |
| 3 | BFI | collective | -190.50 | 391.00 | 0.00 | 1.00 | 1.00 | 0.04 | 0.00 | 2.82 | 0.16 | 0.25 | 5 | 0.09 | 0.33 |
| 4 | BFI | individual | -211.30 | 426.59 | 35.60 | 0.00 | 0.00 | 1.00 | 9.47 | | 0.10 | | 2 | 0.18 | 0.00 |
| 5 | BLO | collective | -221.03 | 452.05 | 0.00 | 1.00 | 1.00 | 0.21 | 1.79 | 0.57 | 0.07 | 0.36 | 5 | 0.07 | 0.76 |
| 6 | BLO | individual | -237.39 | 478.78 | 26.73 | 0.00 | 0.00 | 1.00 | 5505.00 | | 0.00 | | 2 | 0.11 | 0.24 |
| 7 | BUR | collective | -118.30 | 246.61 | 0.00 | 1.00 | 1.00 | 0.11 | 0.00 | 500.00 | 0.15 | 0.03 | 5 | 0.07 | 0.82 |
| 8 | BUR | individual | -143.72 | 291.44 | 44.83 | 0.00 | 0.00 | 1.00 | 10.00 | | 0.10 | | 2 | 0.18 | 0.01 |
| 9 | CAD | collective | -48.28 | 106.56 | 2.65 | 0.27 | 0.21 | 0.17 | 0.00 | 0.43 | 1.46 | 0.39 | 5 | 0.08 | 0.95 |
| 10 | CAD | individual | -49.95 | 103.91 | 0.00 | 1.00 | 0.79 | 1.00 | 0.00 | | 0.39 | | 2 | 0.11 | 0.74 |
| 11 | GIL | collective | -74.54 | 159.08 | 0.00 | 1.00 | 1.00 | 0.26 | 0.00 | 0.00 | 0.33 | 0.75 | 5 | 0.14 | 0.06 |
| 12 | GIL | individual | -102.31 | 208.62 | 49.53 | 0.00 | 0.00 | 1.00 | 2.04 | | 0.35 | | 2 | 0.20 | 0.00 |
| 13 | HER | collective | -136.97 | 283.94 | 2.08 | 0.35 | 0.26 | 0.28 | 0.00 | 0.16 | 0.38 | 0.18 | 5 | 0.08 | 0.95 |
| 14 | HER | individual | -138.93 | 281.86 | 0.00 | 1.00 | 0.74 | 1.00 | 0.00 | | 0.18 | | 2 | 0.11 | 0.59 |
| 15 | HHS | collective | -61.35 | 132.70 | 0.00 | 1.00 | 1.00 | 0.10 | 0.00 | 0.00 | 0.14 | 0.50 | 5 | 0.13 | 0.36 |
| 16 | HHS | individual | -83.75 | 171.50 | 38.80 | 0.00 | 0.00 | 1.00 | 7.10 | | 0.13 | | 2 | 0.24 | 0.01 |
| 17 | HOR | collective | -40.50 | 91.00 | 0.00 | 1.00 | 1.00 | 0.06 | 0.00 | 0.54 | 0.22 | 0.63 | 5 | 0.05 | 0.97 |
| 18 | HOR | individual | -56.65 | 117.30 | 26.30 | 0.00 | 0.00 | 1.00 | 7.77 | | 0.20 | | 2 | 0.17 | 0.02 |
| 19 | MAK | collective | -84.52 | 179.04 | 0.00 | 1.00 | 1.00 | 0.21 | 2.32 | 0.02 | 0.12 | 0.48 | 5 | 0.09 | 0.79 |
| 20 | MAK | individual | -98.02 | 200.05 | 21.01 | 0.00 | 0.00 | 1.00 | 9.17 | | 0.10 | | 2 | 0.19 | 0.03 |
| 21 | MEL | collective | -115.32 | 240.63 | 0.00 | 1.00 | 1.00 | 0.08 | 0.00 | 0.00 | 0.21 | 0.34 | 5 | 0.16 | 0.07 |
| 22 | MEL | individual | -135.73 | 275.46 | 34.83 | 0.00 | 0.00 | 1.00 | 0.00 | | 0.25 | | 2 | 0.24 | 0.00 |
| 23 | MPA | collective | -181.87 | 373.75 | 0.00 | 1.00 | 0.96 | 0.57 | 0.58 | 0.00 | 0.19 | 0.42 | 5 | 0.07 | 0.87 |
| 24 | MPA | individual | -188.04 | 380.08 | 6.33 | 0.04 | 0.04 | 1.00 | 1.47 | | 0.19 | | 2 | 0.10 | 0.43 |
| 25 | ROT | collective | -138.13 | 286.26 | 0.00 | 1.00 | 1.00 | 0.31 | 0.00 | 0.00 | 0.31 | 0.48 | 5 | 0.15 | 0.02 |
| 26 | ROT | individual | -153.36 | 310.71 | 24.45 | 0.00 | 0.00 | 1.00 | 0.00 | | 0.35 | | 2 | 0.20 | 0.00 |
| 27 | SAU | collective | -108.18 | 226.37 | 0.00 | 1.00 | 0.99 | 0.31 | 0.00 | 9.98 | 0.26 | 0.16 | 5 | 0.06 | 0.96 |
| 28 | SAU | individual | -115.50 | 235.01 | 8.64 | 0.01 | 0.01 | 1.00 | 1.49 | | 0.27 | | 2 | 0.13 | 0.21 |
| 29 | SOM | collective | -70.54 | 151.07 | 0.00 | 1.00 | 0.96 | 0.00 | 10.00 | 10.00 | 1.50 | 0.15 | 5 | 0.11 | 0.50 |
| 30 | SOM | individual | -76.67 | 157.34 | 6.27 | 0.04 | 0.04 | 1.00 | 0.00 | | 0.38 | | 2 | 0.16 | 0.10 |
| 31 | SRA | collective | -123.37 | 256.74 | 0.00 | 1.00 | 1.00 | 0.24 | 0.00 | 0.21 | 0.21 | 0.49 | 5 | 0.05 | 0.98 |
| 32 | SRA | individual | -138.11 | 280.23 | 23.49 | 0.00 | 0.00 | 1.00 | 3.00 | | 0.21 | | 2 | 0.14 | 0.07 |
| 33 | STU | collective | -155.70 | 321.40 | 4.32 | 0.12 | 0.10 | 0.35 | 0.00 | 1.02 | 0.52 | 0.16 | 5 | 0.05 | 1.00 |
| 34 | STU | individual | -156.54 | 317.08 | 0.00 | 1.00 | 0.90 | 1.00 | 1.35 | | 0.16 | | 2 | 0.05 | 1.00 |
| 35 | SWP | collective | -40.42 | 90.85 | 0.00 | 1.00 | 0.95 | 0.00 | 0.25 | 1.27 | 0.00 | 0.38 | 5 | 0.10 | 0.71 |
| 36 | SWP | individual | -46.44 | 96.88 | 6.03 | 0.05 | 0.05 | 1.00 | 0.00 | | 0.42 | | 2 | 0.14 | 0.31 |
| 37 | YAL | collective | -205.76 | 421.52 | 0.00 | 1.00 | 1.00 | 0.08 | 0.00 | 0.30 | 0.18 | 0.27 | 5 | 0.04 | 0.99 |
| 38 | YAL | individual | -222.43 | 448.87 | 27.35 | 0.00 | 0.00 | 1.00 | 0.22 | | 0.21 | | 2 | 0.10 | 0.34 |
| 39 | ZSL | collective | -110.01 | 230.02 | 0.00 | 1.00 | 1.00 | 0.24 | 0.00 | 0.79 | 0.39 | 0.50 | 5 | 0.04 | 1.00 |
| 40 | ZSL | individual | -119.95 | 243.90 | 13.88 | 0.00 | 0.00 | 1.00 | 0.39 | | 0.42 | | 2 | 0.10 | 0.25 |

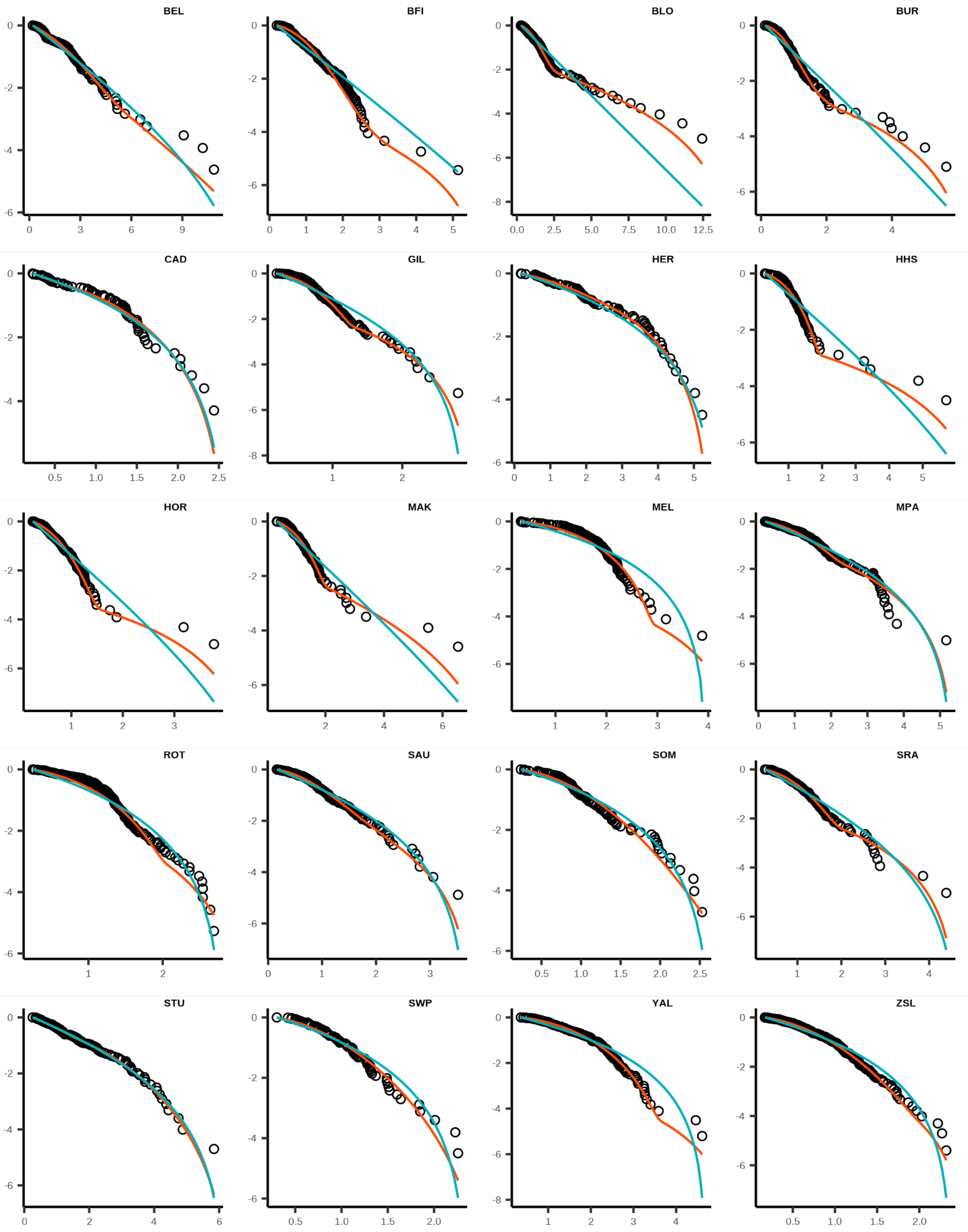


FIGURE S1. **Cumulative frequency distribution** of waggle dance durations and fits of the collective (red line) and individual (blue line) models for all 20 sites

Table S2. Percentage area covered for each land-use type in the agri-rural and urban environments in the sites studied.

| Environment | Land-use | % coverage |
|---|---|---|
| Agri-rural | Arable | 28.30 |
| Agri-rural | Pasture | 23.30 |
| Agri-rural | Woodland | 21.10 |
| Agri-rural | Built Up Area | 15.00 |
| Agri-rural | Fruit | 3.10 |
| Agri-rural | Oilseed Rape | 2.90 |
| Agri-rural | Non Agricultural Unmanaged Green Space | 2.80 |
| Agri-rural | Non Agricultural Managed Green Space | 1.80 |
| Agri-rural | Other Agricultural | 1.50 |
| Agri-rural | Water | 0.20 |
| Urban | Sparse Residential | 34.80 |
| Urban | Continuous Central | 24.30 |
| Urban | Dense Residential | 21.80 |
| Urban | Parks Allotments Cemeteries | 7.90 |
| Urban | Woodland | 4.20 |
| Urban | Water | 3.40 |
| Urban | Amenity Grassland | 2.60 |
| Urban | Railway | 1.00 |

### Jackknifed partial least squares analysis

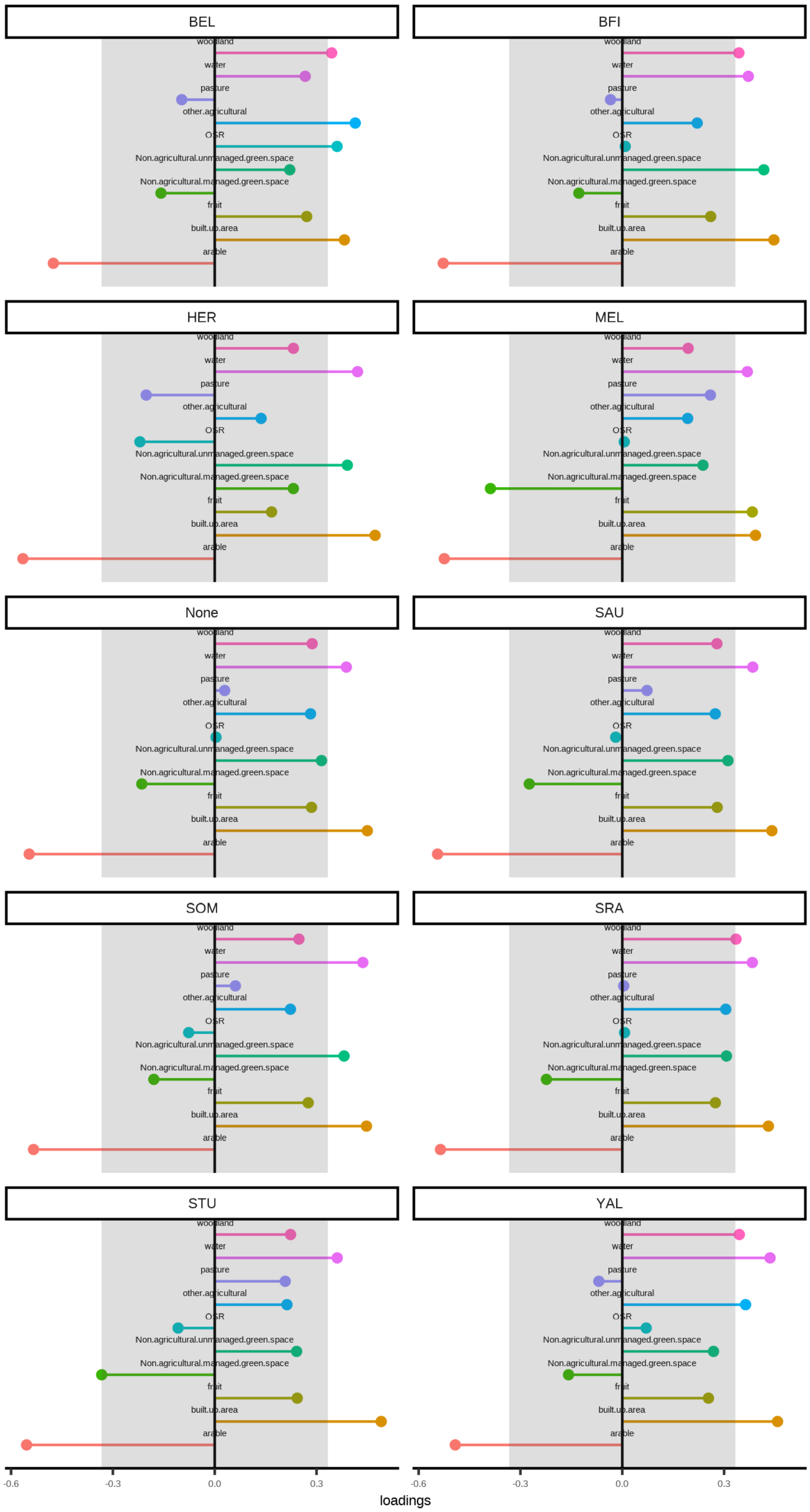


Figure S2. **Loadings of PLS calculated for each site removed for the agri-rural sites**. Each plot shows the loadings of the first principle component with that site removed from the analysis, showing the individual points making up the overall box plot loadings in Fig 4b.
